# 3D morphometric analysis of mouse skulls using microcomputed tomography and computer vision

**DOI:** 10.1101/2022.10.26.513830

**Authors:** Beatrice R. Gulner, Zahra S. Navabi, Suhasa B. Kodandaramaiah

**Affiliations:** Department of Mechanical Engineering, University of Minnesota Twin Cities, Minneapolis, MN, USA; Department of Biomedical Engineering, University of Minnesota Twin Cities, Minneapolis, MN, USA

## Abstract

Morphometric studies have provided crucial insights into the skull anatomy of commonly used wildtype (WT) laboratory mice strains such as the C57BL/6. With the increasing use of transgenic (TG) animals in neuroscience research, it is important to determine whether the results from morphometric studies performed on WT strains can be extended to TG strains derived from these WT strains. We report a new computer vision-based analysis pipeline for surveying mouse skull morphology using microcomputed tomography (μCT) scans. We applied this pipeline to study and compare eight cohorts of adult mice from two strains, including both male and female mice at two age points. We found that the overall skull morphology was generally conserved between cohorts, with only 13% of landmark distance differences reaching statistical significance. In addition, we surveyed the dorsal skull bone thickness differences between cohorts. We observed significantly thicker dorsal, parietal, and/or interparietal bones in WT, male, or older mice for 53% of thickness comparisons. This knowledge of dorsal skull bone thickness has potential implications for surgical planning through skull imaging and has applications in automating cranial microsurgeries on mice.

## Introduction

Mice (mus musculus), with their small size, relatively short breeding and developmental cycle and well conserved brain morphology have emerged as one of the most widely used mammalian model organisms in neuroscience^1^. Recently developed strategies for cell type-specific expression of genetically encoded neural activity reporters and perturbations^2,3^ have facilitated creation of a wide range of new TG mouse strains.

Targeting specific brain regions in mice for virus injections, insertion of penetrating neural interfaces or implantation of cranial windows for imaging generally relies on skull landmark identification and measurement during stereotactic surgery. A previous morphometric study of postnatal skull ontogeny^4^ used μCT scans to show that after male C57BL/6 mice reach adulthood, the growth in overall shape and size of the skull plateaus. Thus, current approaches for stereotactic targeting are highly reliable and accurate. However, current morphometrics studies focus primarily on common WT strains^5^. Generation of TG strains can result in unintended phenotypic changes^6^. Not much is known about similarities between TG mice skull morphology and the WT strains they are derived from. Further, a detailed understanding of the variation of the thickness of bone in the skull is not known. Knowledge of skull bone thickness would allow better planning of cranial surgeries.

Here, we combine μCT scanning of mouse skulls with a computer vision analysis pipeline to perform morphometric analyses on an in-house bred Thy1-GCaMP6f TG mice strain^7^. These results were then compared with WT C57BL/6 mice of comparable ages obtained from a commercial vendor. We also investigated the effect of sex and age on the skull morphometrics of both strains. Two aspects of skull morphology were considered: the sizes of representative features in the region surrounding the cranial cavity and the thickness of the bone of the dorsal skull. While traditional morphometric techniques can measure externally accessible skull features, measurement of dorsal skull bone thickness from intact specimens requires a non-destructive imaging technique such as μCT.

## RESULTS

### Experimental and image analysis workflow

Skull specimens were preserved in 4% paraformaldehyde and scanned in a μCT x-ray scanner (**Fig. 1A**). μCT scans were reconstructed and registered to a common reference frame using commercial software packages CT Pro 3D and VGStudio MAX 3.2, then imported into MATLAB as a coronal section image stack (**Fig. 1B**). Bone was segmented from background using an Otsu’s method-based threshold on the grayscale intensity values (**Fig. 1C**). Distances between eight pairs of morphological landmarks were used to characterize the shape of the dorsal skull and cranial cavity (**Fig. 2A**). The images after thresholding were analyzed using custom MATLAB scripts which measured the thickness of the bone across the dorsal skull (**Fig. 1D**). For each cohort, scans were co-registered and averaged elementwise to create a representative skull (**Supplementary Fig. 1A-C**). Average thickness was calculated by dorsal skull bone region of interest (ROI) for each cohort representative skull, then Students’ t-tests evaluated statistical significance of thickness differences between cohorts (**Supplementary Fig. 2A-D**).

**Figure 1.**
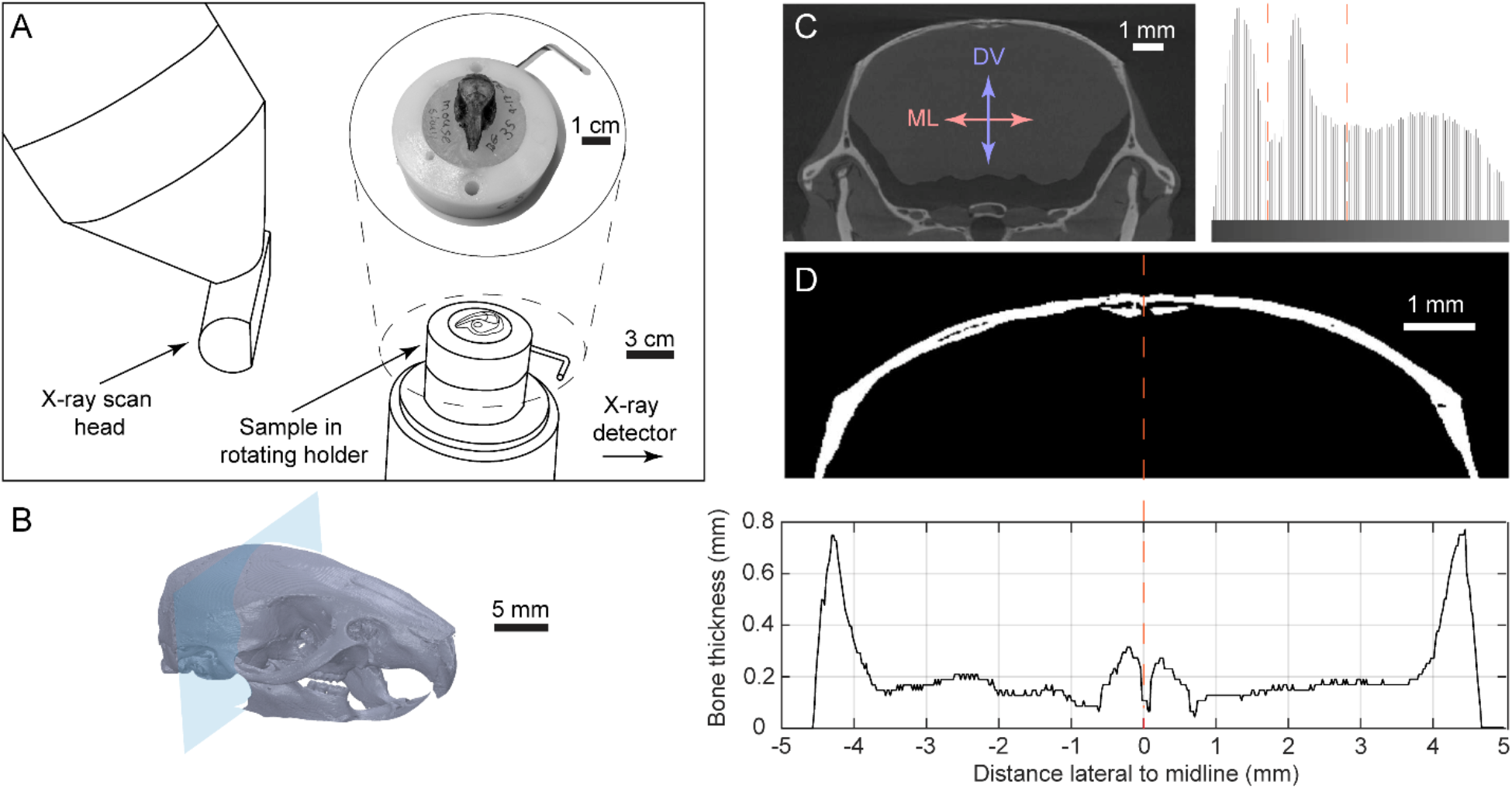
Microcomputed Tomography (μCT) scanning and image stack measurement: **(A)** μCT scanner setup and example mounted sample. **(B)** Reconstructed three-dimensional scan of skull with coronal section indicated by intersecting plane. **(C)** Raw cranial section from μCT scan image stack and histogram of grayscale intensity values. Dashed lines indicate segmentation thresholds identified using a modified Otsu’s Method. **(D)** Segmented image cropped to area of interest and corresponding plot of skull bone thickness, defined as the distance in millimeters between the first and last white pixels in each column of the image. Plot trace matches above image and dashed line indicates midline location.

**Figure 2.**
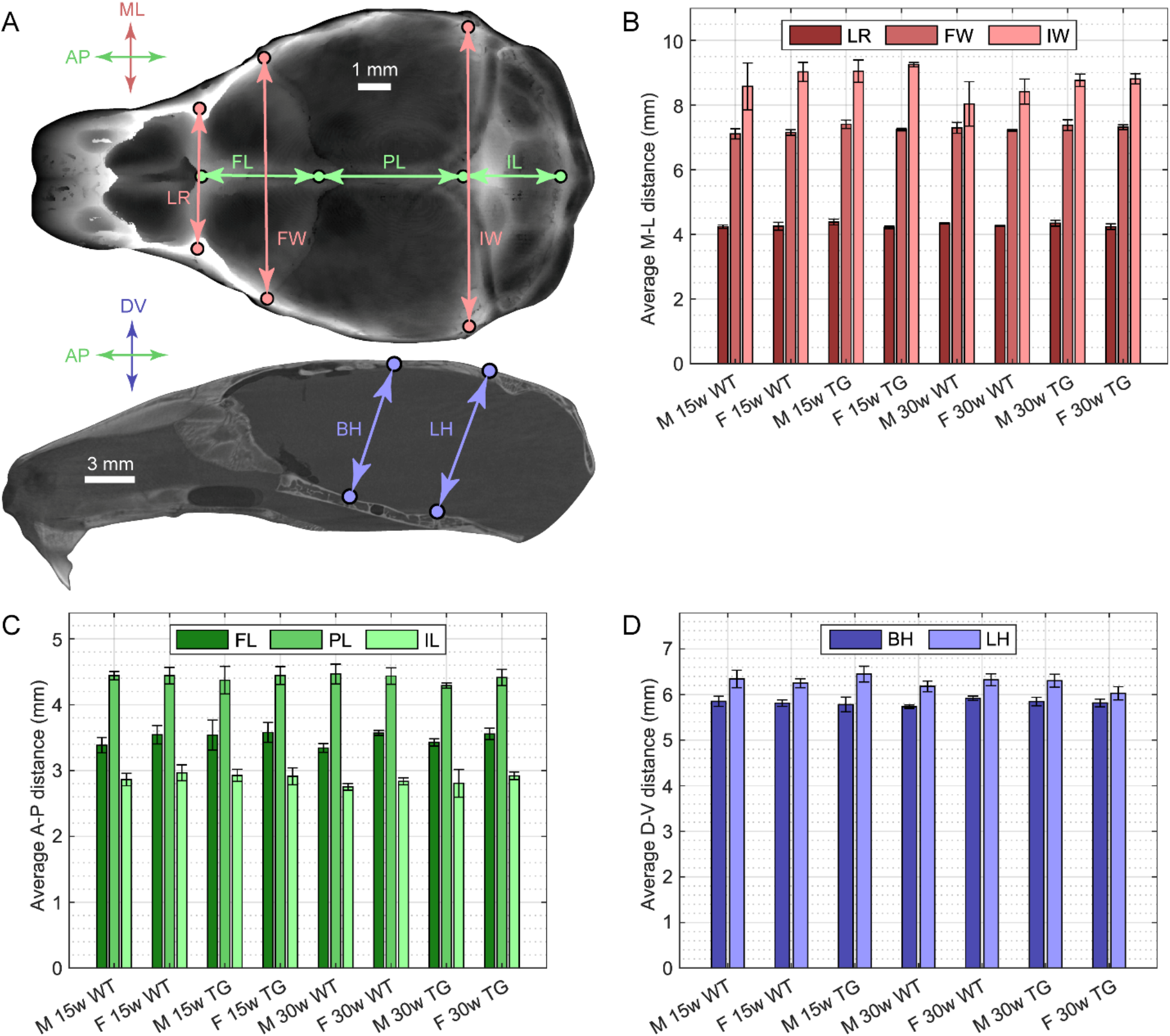
Morphological landmark distances: **(A)** Distance measurements and landmarks indicated on 2D plot of dorsal skull thickness measurements (top) and a sagittal section at Bregma (bottom). Distance measurements are abbreviated as follows: LR – width between anterolateral corners of the frontal bone; FW – width of the frontal bone measured at the lateral ends of the coronal suture; FL – length of the frontal bone measured parallel to the midline; PL – length of the parietal bone measured parallel to the midline; IW – width of the interparietal bone measured at the lateral ends of the lambdoid suture; IL – length of the interparietal bone measured parallel to the midline; BH – height of the cranial cavity measured between Bregma and the intersphenoidal synchondrosis; LH – height of the cranial cavity measured between Lambda and the sphenooccipital synchondrosis. **(B-D)** Bar plots showing the average of the measured distances between landmark pairs divided by cohort; **(B)** medial-lateral (M-L) direction; **(C)** anterior-posterior (A-P) direction; **(D)** dorsal-ventral (D-V) direction. Error bars indicate one standard deviation.

### Comparison with published morphological landmark distances

The distance between landmarks on the skull quantifies skull shape morphology. To validate our measurement methodology, we compared the distances measured in our study with published data from studies which investigated 16-week C57BL/6 male mice^4^ or 12-week C57BL/6 male and female mice^5^. We measured C57BL/6 mice at the age of 15-weeks (n=5 male and n=5 female). Six distances (**Fig. 2A**) with analogues in the literature were identified: FW – width of the frontal bone measured at the lateral ends of the coronal suture; PL – length of the parietal bone; IW – width of the interparietal bone measured at the lateral ends of the lambdoid suture; IL – length of the interparietal bone; BH – height of the cranial cavity between Bregma and the intersphenoidal synchondrosis; LH – height of the cranial cavity between Lambda and the spheno-occipital synch-ondrosis. All lengths were measured parallel to the midline. The comparisons are summarized in **Table 1**.

**Table 1:**
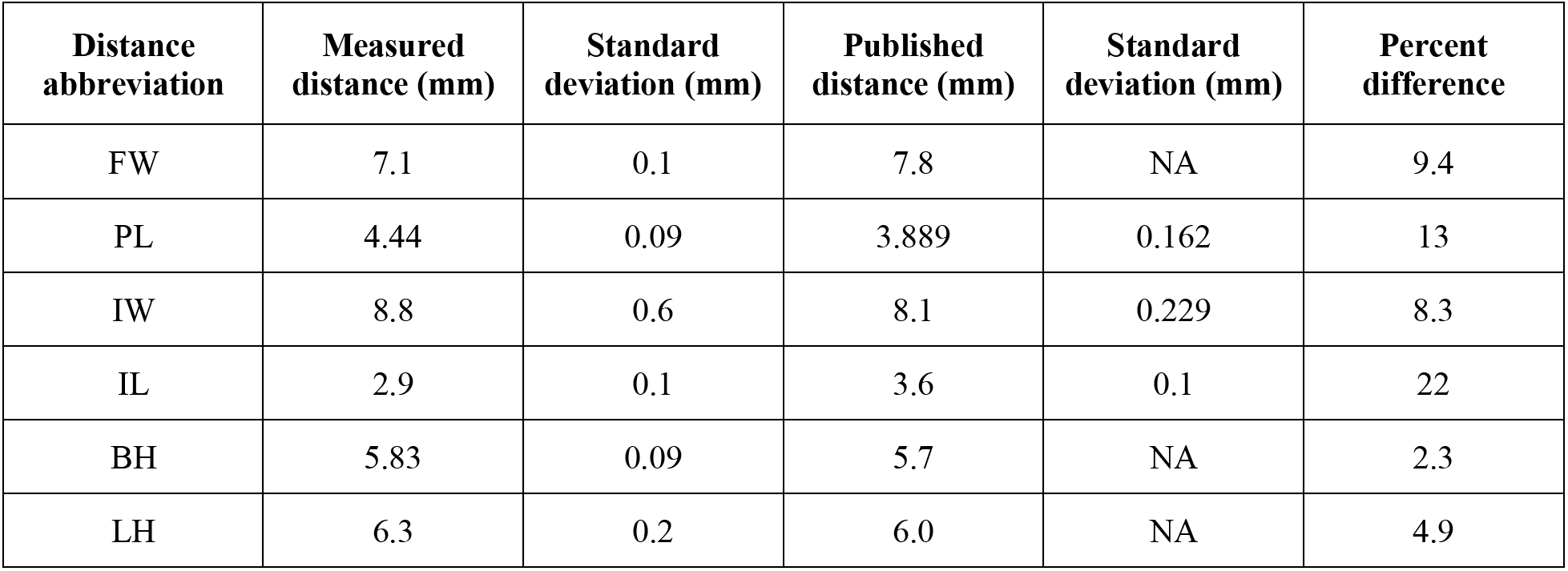
Distances between landmarks compared with published values^4,5^. Standard deviations are provided when available. The distance measurements are abbreviated as follows (see **Fig. 2A**): FW – width of the frontal bone measured at the lateral ends of the coronal suture; PL – length of the parietal bone; IW – width of the interparietal bone measured at the lateral ends of the lambdoid suture; IL – length of the interparietal bone measured; BH – height of the cranial cavity between Bregma and the intersphenoidal synchondrosis; LH – height of the cranial cavity between Lambda and the spheno-occipital synchondrosis. All lengths are measured parallel to the midline. Published data provided comparison values for FW, BH, LH^4^ and PL, IW, IL^5^ respectively.

| Distance abbreviation | Measured distance (mm) | Standard deviation (mm) | Published distance (mm) | Standard deviation (mm) | Percent difference |
| --- | --- | --- | --- | --- | --- |
| FW | 7.1 | 0.1 | 7.8 | NA | 9.4 |
| PL | 4.44 | 0.09 | 3.889 | 0.162 | 13 |
| IW | 8.8 | 0.6 | 8.1 | 0.229 | 8.3 |
| IL | 2.9 | 0.1 | 3.6 | 0.1 | 22 |
| BH | 5.83 | 0.09 | 5.7 | NA | 2.3 |
| LH | 6.3 | 0.2 | 6.0 | NA | 4.9 |

In general, our measurements agree well with both sets of published results^4,5^. The largest deviation was the IL length of (2.9 ± 0.1) mm versus the published (3.6 ± 0.1) mm, a percent difference of 22%. The dorsal-ventral (D-V) height measurements had the lowest percent differences, with 2.3% and 4.8% for BH and LH respectively. The mean percent difference was 10% across all measurements. The differences may be due to natural variation between mice or inherent inconsistencies in manual landmark identification. Since the magnitudes of the percent differences are relatively small, using μCT scans and our computer vision analysis pipeline to measure morphological landmark distances is reasonable.

### Differences in morphological landmark distances between cohorts

We next compared the landmark distances in TG Thyl-GCaMP6f mice^7^ with corresponding values for the WT C57BL/6 strain. The Thy1-GCaMP6f mice were derived from the C57BL/6 line and were bred within our in-house colony. These comparisons thus evaluated any differences between WT and TG mice, while also accounting for differences between in-house bred versus commercially procured mice. Based on prior studies, we expected that morphology would be relatively constant for adult mice, so minimal differences were expected between age points^4^.

In addition to the six landmark distances described in the previous section, we included LR – width between the anterolateral corners of the frontal bone; and FL – length of the frontal bone measured parallel to the midline. Comparisons were made for each distance between pairs of cohorts where one variable changed and the other two were held constant. For example, the cohort of 15-week male WT mice was compared with the cohort of 30-week male WT mice, varying the age variable while holding sex and strain constant. 13 of the 90 distance comparisons yielded significant differences (Students t-test). **Figures 2B-D** show the landmark distance measurements. The significance results for each comparison are reported in **Table 2**.

**Table 2:**
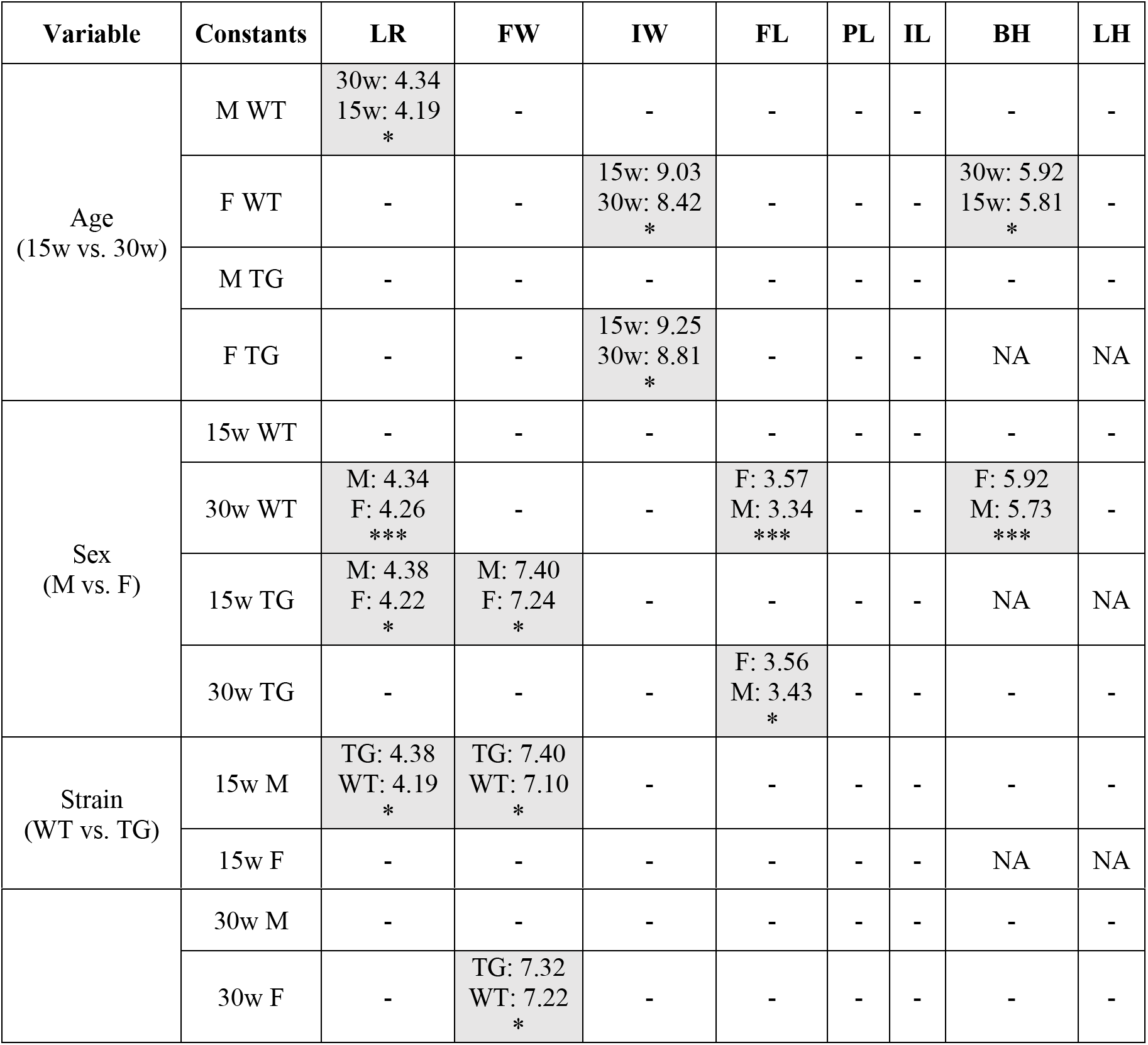
Results from the landmark distance comparison between cohorts. Abbreviations added to those from **Table 1**: LR – width between the anterolateral corners of the frontal bone; FL – length of the frontal bone measured parallel to the midline. Distances given in mm. * indicates p < 0.05; *** indicates p < 0.001, Students’ t-test. Dashes indicate no significant differences. “NA” indicates distance could not be measured.

None of the measurable PL, IL, or LH comparisons were significant. Nor were any comparisons between cohorts with differing ages for male TG mice, sexes for 15-week WT mice, strains for 15-week female mice, or strains for 30-week male mice. Between strains, only the medial-lateral (M-L) width comparisons had significant differences, all comparisons indicating wider skulls in TG mice. For the six significant sex comparisons, the male skulls were wider in the M-L direction while the female skulls were longer in the anterior-posterior (A-P) direction and taller in the D-V direction. Only M-L width and D-V height showed any significant differences in age comparisons.

### Dorsal skull bone thickness analysis

Most neuroscience studies requiring invasive or minimally invasive neural recording and manipulation involve performing small to large craniotomies in the skull. For a successful craniotomy procedure, the bone must be removed completely, efficiently, and with little or no damage to the underlying soft tissues like dura and brain. Efforts have been made in recent years to automate this procedure using impedance sensing feedback^8^ or force feedback^9,10^. These studies have found that there is substantial variation in the thickness of the dorsal skull, both between subregions of the skull for a single mouse and between different mice at the same location on the skulls. In this study, we used the μCT scan database to systematically evaluate the variation in skull thickness across cohorts of mice.

We constructed representative half skulls for each cohort. A custom control point registration algorithm employing piecewise linear transformations aligned each half skull to a set of reference points. The sets of registered skulls were then averaged elementwise to create the representative skulls for each cohort (**Supplementary Fig. 1A-C**).

An example comparing an individual half skull from the cohort of female 30-week WT mice with the cohort average skull is shown in **Figure 3A**. The qualitative similarities between the features on the individual and cohort average skulls indicate that the registration was successful and suggest that the finer patterns of bone thickness were consistent between mice once the gross features, such as sutures, sinuses, and peripheral cranial cavity edges, were aligned.

**Figure 3.**
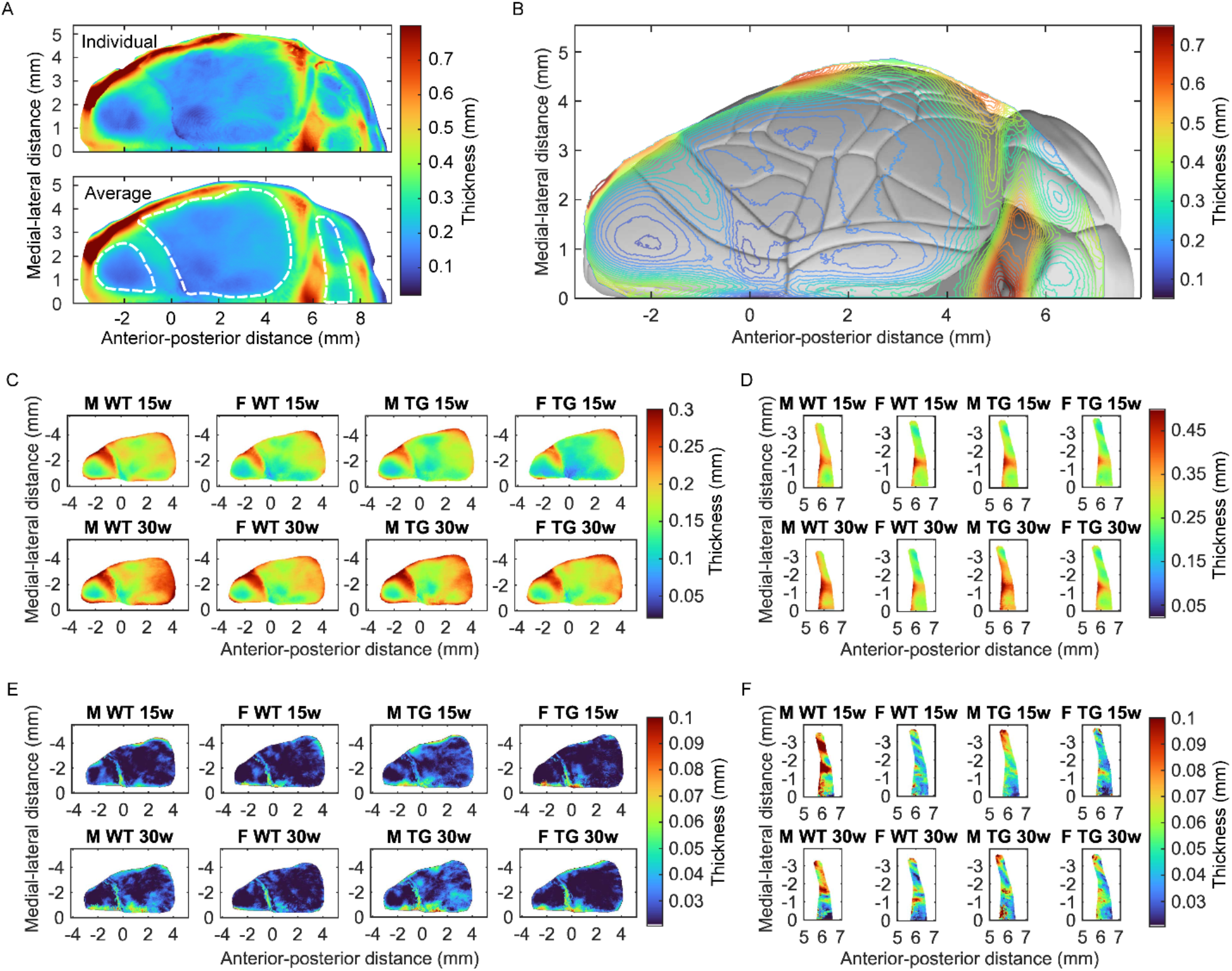
Bone thickness profile across dorsal skull: **(A)** Comparison between the individual skull from the female 30-week wildtype (WT) cohort (top) and the corresponding cohort average skull (bottom). Similarity between gross structures on the skulls indicates successful registration, while consistency in the finer features suggests uniformity between skulls on the scale of bone subregions. Region of interest (ROI) boundaries indicated by dashed white lines superimposed on cohort average plot. From anterior to posterior: frontal ROI, parietal ROI, and interparietal ROI. **(B)** Contour plot of the average skull from all scans in the study superimposed on the Allen Mouse Brain Connectivity Atlas, connectivity.brain-map.org/. Patterns of subregions with consistent thickness align with anatomically distinct regions of the cerebellum and midbrain, as expected, but also with functional regions on the lissencephalic cortex. **(C-D)** Average skulls for each cohort, frontal and parietal **(C)** and interparietal **(D)** ROIs. There is apparent similarity in the bone thickness profile between cohorts. Note the larger range in thickness color bar values required for interparietal ROI pseudocolor plots. **(E-F)** Standard deviation of average skulls by cohort, frontal and parietal **(E)** and interparietal **(F)** ROIs. Low standard deviations suggest minimal variation in bone thickness profile within cohorts after registration. Generally larger standard deviation values for interparietal than for frontal and parietal ROIs.

Qualitatively, we observed consistent patterns of skull thickness variations across the dorsal skull, particularly in the frontal and parietal bones. There were distinct thinner sections separated by thicker ridges in the interparietal bone, which envelopes the cerebellum and midbrain. The thinner sections loosely overlapped with anatomically distinct subregions, including the declive, culmen, simple lobule, and ansiform lobules in the cerebellum as well as the superior and inferior colliculus in the midbrain.

In contrast, the dorsal cortex in the mouse brain is lissencephalic. A contour plot of the representative half skull comprising all mice in the study was superimposed on the Allen Brain Atlas (**Fig. 3B**). A rough correspondence between regions of uniform skull thickness and functional regions of the cortex was apparent, including for the frontal and parietal bones. For example, there are relatively thinner sections of the parietal bone roughly aligned above the retrosplenial cortex and the primary somatosensory area barrel field level 2/3. There is also a relatively thinner section of frontal bone above the secondary motor area and a thicker section above the primary motor area. These qualitative correlations perhaps indicate a closer relationship between the skull and functionally distinct regions of the dorsal cortex.

Similar patterns of bone thickness were observed across the dorsal skull in all cohorts, though the thickness contours appeared to scale by the overall thickness magnitude of the cohort (**Fig. 3C,D**). As an example, again consider the average skull for the female 30-week WT cohort. The average thickness differs substantially between the frontal/parietal and interparietal ROIs, with an average thickness of 0.19 ± 0.04 mm and 0.28 ± 0.06 mm respectively. The maximum thickness for the frontal/parietal region is 0.33 mm, found 2.9 mm lateral and 1.7 mm anterior to Bregma, on the lateral edge of the coronal suture. The minimum thickness is 0.086 mm, located 0.62 mm directly lateral to Bregma. The maximum thickness for the interparietal ROI is 0.49 mm, found 1.4 mm lateral and 5.8 mm posterior to Bregma, or just posterior to the lambdoid suture. The minimum thickness of 0.17 mm is located 3.0 mm lateral and 6.0 mm posterior to Bregma, or slightly posterior to lambda and far lateral, above the ansiform lobule.

Plots of the elementwise standard deviation for each cohort representative skull indicate minimal variation in dorsal skull thickness within cohorts (**Fig. 3E,F**). The maximum variance for the female 30-week WT frontal/parietal ROI was located 0.64 mm lateral and 3.0 mm anterior to Bregma, likely due to proximity to the midline. The least variance was found 2.7 mm lateral and 2.0 mm posterior to Bregma. The thickness of central regions of the parietal bone generally appears more uniform. In the interparietal ROI, the maximum variance in skull thickness was located 0.50 mm lateral and 5.8 mm posterior to Bregma, likely due to proximity to the lambdoid suture. The minimum standard deviation is found 0.042 mm lateral and 6.6 mm posterior to Bregma, on the far posterior edge of the ROI. Overall, the standard deviations are small, with maxima tending to fall near sutures where the registration imperfectly aligned the fine features.

The average thickness across each of the frontal, parietal, and interparietal ROIs was computed by skull and compiled into vectors by cohort (**Supplementary Fig. 2A-D**). Tests of statistical significance were applied to the average thickness vectors for every pair of cohorts which differed by only one variable (Students’ two-sample t-test, p < 0.05). Twelve pairs of cohorts were compared for each ROI. The null hypothesis was that the cohorts had equal average bone thicknesses across the ROI; the alternative hypothesis was that the averages differed. **Figure 4A** shows the measurements of average bone thickness. Box plots summarized each significance test; see example in **Figure 4B**. The significance results for each comparison are reported in **Table 3**.

**Figure 4.**
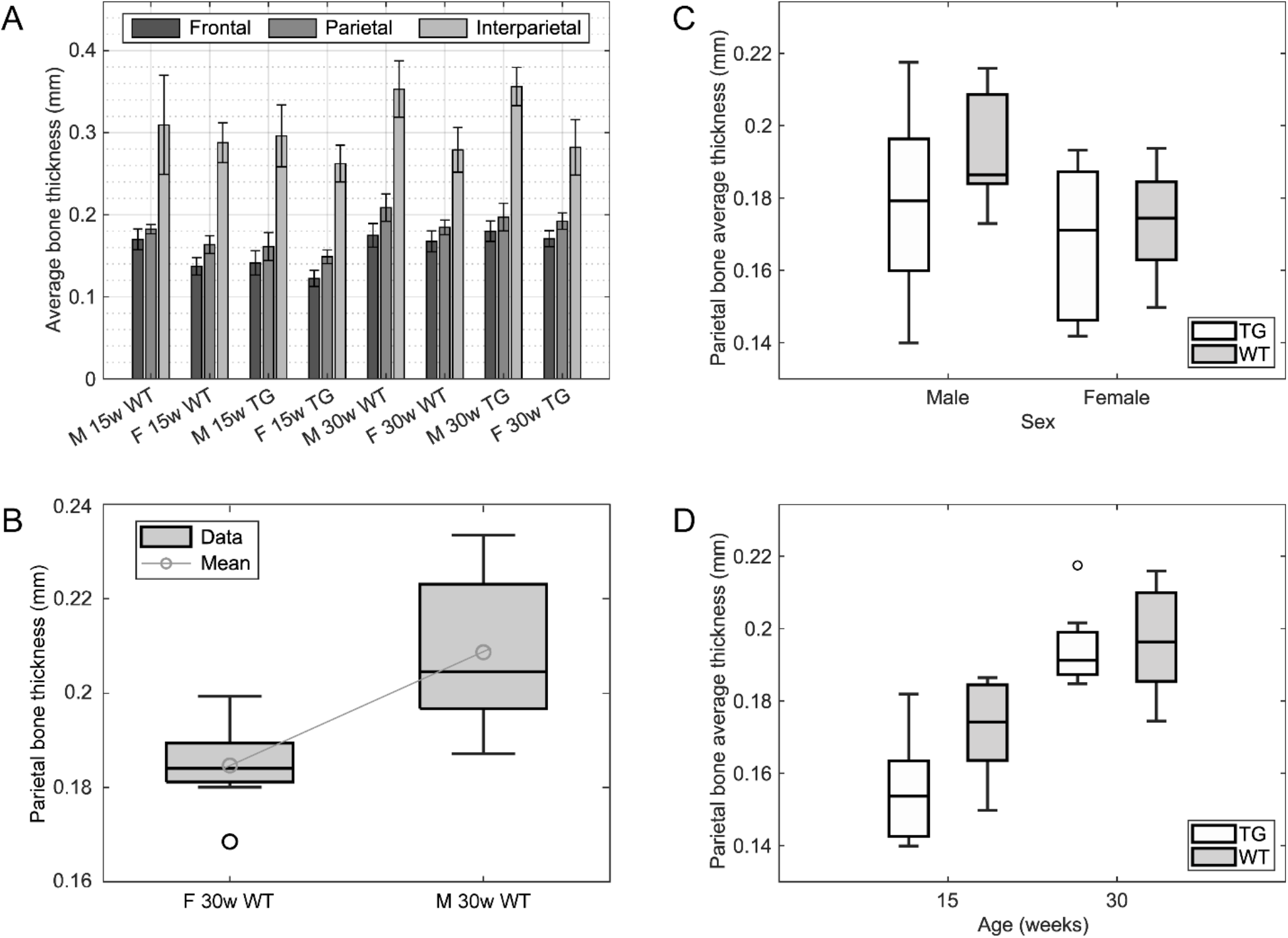
Average dorsal skull bone thickness differences between cohorts: **(A)** Average bone thickness across frontal, parietal, and interparietal ROIs, divided by cohort. Error bars indicate one standard deviation. **(B)** Example of box-and-whisker plots which summarize the mean and distribution of dorsal skull bone thickness for each comparison between cohorts. **(C)** Box-and-whisker plot of average parietal ROI thickness divided by strain with sex as the independent variable. **(D)** Box-and-whisker plot of average parietal ROI thickness divided by strain with age as the independent variable.

**Table 3:** Results from the comparisons of average dorsal skull bone thickness between cohorts. Thicknesses given in mm. * indicates p < 0.05; *** indicates p < 0.001 (t-test). Dashes indicate no significant differences.

| Variable | Constants | Frontal | Parietal | Interparietal |
| --- | --- | --- | --- | --- |
| Age<br>(15w vs. 30w) | M WT | - | 30w: 0.21<br>15w: 0.18<br>*** | - |
|  | F WT | 30w: 0.17<br>15w: 0.14<br>*** | 30w: 0.18<br>15w: 0.16<br>*** | - |
|  | M TG | 30w: 0.18<br>15w: 0.14<br>*** | 30w: 0.20<br>15w: 0.16<br>*** | 30w: 0.36<br>15w: 0.30<br>*** |
|  | F TG | 30w: 0.17<br>15w: 0.12<br>*** | 30w: 0.19<br>15w: 0.15<br>*** | - |
| Sex<br>(M vs. F) | 15w WT | M: 0.17<br>F: 0.14<br>*** | M: 0.18<br>F: 0.16<br>*** | - |
|  | 30w WT | - | M: 0.21<br>F: 0.18<br>* | M: 0.35<br>F: 0.28<br>*** |
|  | 15w TG | M: 0.14<br>F: 0.12<br>* | - | M: 0.30<br>F: 0.26<br>* |
|  | 30w TG | - | - | M: 0.36<br>F: 0.28<br>*** |
| Strain<br>(WT vs. TG) | 15w M | WT: 0.17<br>TG: 0.14<br>*** | WT: 0.18<br>TG: 0.16<br>*** | - |
|  | 15w F | WT: 0.14<br>TG: 0.12<br>* | WT: 0.16<br>TG: 0.15<br>* | - |
|  | 30w M | - | - | - |
|  | 30w F | - | - | - |

19 of 36 total comparisons were significant. The dorsal skull bones of 30-week mice were thicker than for 15-week mice in at least one comparison for every ROI, including all parietal ROI comparisons. Male mice had higher average bone thickness than female mice in at least one comparison for every ROI. WT mice had thicker dorsal skull bones than TG mice in all 15-week frontal and parietal ROI comparisons. There were no significant differences between strains for 30-week cohorts or the interparietal ROI.

We examined bone thickness variation by strain when the cohorts were additionally separated by sex or age, results shown in **Figures 4C** and **D** respectively. Only the parietal bone ROI was considered as its boundaries were most reliably identifiable. The parietal ROI thickness was comparable between WT and TG strains for both male and female mice. The WT parietal bones were thicker at 15 weeks, but the thickness equalized at 30 weeks. Perhaps the WT parietal bones thicken faster than the TG, but by 30 weeks the TG mice development has caught up.

## DISCUSSION

The minimal measurable differences in the morphometry of mouse skulls across strains indicates that the current approach of using brain atlas maps of WT C57BL/6 mice^11,12^ as a ground truth for planning stereotactic surgeries should continue to be effective in Thy1-GCaMP6f and other TG strains derived from C57BL/6. Further, the few significant differences in age point comparisons indicate that as expected, the overall skull shape remains largely stable once mice reach adulthood.

Across the frontal, parietal, and interparietal bone ROIs which were analyzed for bone thickness, we found that the thickness of the dorsal skull bone varied between certain cohorts. The older mice at 30-weeks had consistently thicker skull bones in all ROIs where significance was achieved, including all parietal ROI comparisons. Male mice also had significantly thicker skull bones than female mice when the comparison was significant, though fewer comparisons reached significance. In the strain comparisons, the WT mice tended to have thicker skull bones than the TG mice, but significance was only reached in comparisons between 15-week mice for the frontal and parietal ROIs. None of the 30-week or interparietal ROI comparisons between strains were significant. Overall, the trends for which demographic had the thicker skull were consistent across comparisons, but not all comparisons achieved statistical significance.

The thickness of the dorsal skull bones of mice varies, both between locations on the skull and between mice with different age, sex, or strain characteristics. In practice, surgeons performing craniotomies must be aware of this non-uniform thickness. They can expect that an older mouse will likely have thicker dorsal skull bones than an otherwise comparable young mouse. It is also probable that a male mouse will have a thicker skull than a female mouse and that a C57BL/6 mouse will have a thicker skull than a Thy1-GCaMP6f mouse. Differences are more likely for the parietal bone than the frontal or especially the interparietal. While we observed that few distances between landmarks differed significantly, the dorsal skull bone thickness did vary with certain changes in mouse characteristics. The thickness thus may vary more than the overall shape morphology of the dorsal skull, at least within practically relevant orders of magnitude. In future, this data could be incorporated into automated cranial surgery platforms^8-10^ to digitally limit the maximum drilling depth.

We note that our results are limited by the 21 μm spatial resolution of the instrumentation. We also studied a single transgenic line. Further study is necessary to determine whether the results hold for other TG lines derived from the same C57BL/6 line.

The ability to image brain structure and physiology through the skull has important applications in neuroscience, particularly in studies where immune disruption caused by implantation of cranial windows^13,14^ is undesirable. Methods for imaging the brain through the skull include using NIR light^15^, visible light^16^, optical coherence tomography^17^, or three-photon imaging^18^. Other approaches include thinned skull imaging and polished skull techniques^19^. Our results indicate that bone thickness variations, which increase light scattering and absorption, may influence the quality of images acquired. This is an experimental variable that should be considered. When using intact skull methods in older animals, the gradual thickening of the skull can also result in lower quality imaging^20,21^. As a final note, there has been increased attention paid to skull microvasculature environment^22^ and the interaction with the meninges. Our results demonstrating increased skull thickness with age indicate possible age-related effects on these interactions.

## METHODS

### Sample preparation

We scanned eight cohorts of four to six mouse skulls each. The cohorts included 19 male and 17 female mice. 18 mice were in-house bred WT C57BL/6 mice and 18 were TG Thy1-GCaMP6f mice from a commercial vendor (Jackson Laboratories). 20 mice were (15 ± 1) weeks old and 16 were (30 ± 1) weeks old. All animal experiments were conducted in accordance with approved University of Minnesota Institutional Animal Care and Use Committee protocol.

Mice were euthanized via isoflurane (Piramal Critical Care Inc., Bethlehem, PA) overdose. The skulls were separated from the cadavers and soft tissue was removed from external surfaces. They were immersed in 4% paraformaldehyde (PFA, CAT# P6148-500G, Sigma Aldrich) for at least 12 hours and stored in a refrigerator.

The skulls were removed from the PFA and rinsed with deionized water. Dental acrylic powder (Dentsply Caulk Orthodontic Resin, York, PA, USA) was mixed with the corresponding curing liquid to form a viscous paste and poured into a mounting ring. The dental acrylic cured for 15-30 seconds before the skull was pressed into the surface of the acrylic. Molding the acrylic by hand as it cured ensured proper orientation of the skull and positioning above the top surface of the mounting ring.

### μCT scanning and reconstruction

Scans were performed using a 225 kV reflection target μCT machine (XT H 225, Nikon Metrology Inc., Brighton, MI, USA). The X-ray settings for all scans were 110 or 120 kV, 85 μA. Each scan consisted of 720 projections at a half degree pitch and took 4 frames for each projection. The exposure time was 708 msec. No filters were used in all but six scans. The location of the skull relative to the scanning head was constant to ± 2 mm.

A 0.5 mm aluminum filter was added in six scans to mitigate edge artifacts. No effect on the results was observed when comparing scans of the same skull with and without the filter, so it was removed for the remaining scans. When present, the artifact is isolated to the anterior medial skull and does not substantially affect the analyses reported here.

The skull scan was reconstructed using a commercial CT reconstruction package (CT Pro 3D, Nikon Metrology Inc., Brighton, MI, USA). We used the simple registration function in VGStudio MAX 3.2 (Volume Graphics GmbH, Heidelberg, Germany) to align the axes of the skull with the scan axes. The midline was aligned with the Z-axis in the transverse plane, the line secant to Bregma and Lambda parallel with the Z-axis in the sagittal plane, and the craniocaudal axis parallel with the Y-axis in the coronal plane. The scan was exported as a coronal section image stack of full-quality JPEG files with the registration preserved but no filtering.

### Segmenting and measuring the dorsal skull bone thickness from the μCT scans

Analysis of the image stacks was performed with custom scripts written in MATLAB (MATLAB R2022a, The MathWorks, Inc., Natick, MA, USA). Parameters were entered manually for each scan, including the image index of the coronal section corresponding to Bregma, the pixel index of the M-L midline location, and the cropping bounds for isolating the dorsal skull. The ventral bound was refined using a coarsely sampled preview of the measured skull to ensure consistent transverse cropping.

Each coronal section image in the scan was segmented using a modified Otsu’s Method algorithm^23^. For consistency across the image stack, the threshold value was determined based on the coronal section at Bregma. The threshold was calculated by averaging the second threshold values from the two- and three-level Otsu threshold results. This balanced optimizing the distinction of bone from background with minimizing edge artifacts. **Figure 1C** shows an unfiltered coronal section and the corresponding histogram with Otsu thresholds indicated, where the rightmost threshold distinguishes bone from background.

Bone thickness was then measured. For each D-V column of pixels, the furthest dorsal and ventral pixels identified as bone were found. The skull thickness at that location was defined as the difference between the pixel indices. The factor 0.021 mm/pixel converted the thickness to millimeters. Comparing the measured profile with the corresponding segmented image confirmed plausibility (**Fig. 1D**). The procedure was repeated for every image in the stack and the bone thickness was displayed in a pseudocolor plot with resolution in each direction of 0.021 mm.

### Computer vision-based co-registration to create an average skull for each cohort

We created an average skull for each cohort using control point registration. A set of control points were identified across the dorsal skull (**Supplementary Fig. 1B**) and manually selected using the MATLAB “drawpoint” function. The furthest anterior control point was used only during registration, not for the landmark distance analysis, as it was added solely to prevent skewing of the anterior region during registration.

Assuming that the skulls are approximately symmetric, we reflected the left halves of the skull across the midline. Reflecting over the midline could increase uncertainty in the results if the midline location was incorrect or if there were lateralized differences between skull halves, but at the resolution of the study is unlikely to cause substantial error. The reflected control points were averaged with the right-side points to eliminate redundant pairs. The unpaired control points along the midline were corrected to lie exactly on the midline.

The A-P and M-L coordinates of corresponding control points were averaged by cohort. We registered each skull in the cohort to this reference set of control points. The “fitgeotrans” function from the MATLAB Image Processing Toolbox was used to estimate the piecewise linear transformation to fit each skull’s control point distribution to the reference set. The “imwarp” function applied the transformation to each matrix of thickness values. After transforming the matrix, the “imwarp” algorithm then interpolated between the transformed points to recreate an evenly spaced grid. The thickness values were averaged elementwise across the entire cohort. The registration process is illustrated in **Supplementary Figure 1A,** and an example of the results is shown in **Supplementary Figure 1C**. We excluded three half-skulls from the analysis which were disturbed during validation of the μCT measurements.

ROIs were selected on the frontal, parietal, and interparietal bones (**Fig. 3A**). Since the measurements were taken parallel to the D-V axis rather than normal to the skull surface, the outer edges of the scans appeared artificially thick and thus were excluded. We also excluded the sinuses and sutures. Cropping out these areas permitted a narrower color bar range for the bone thickness pseudocolor plots, thus revealing more of the fine bone structure. Most cranial surgeries are performed on the central bone regions, so data focusing on these areas should be sufficient for many applications. We registered the ROI masks to each skull using the control point registration method.

The preparatory steps for the statistical analysis are illustrated in **Supplementary Figures 2A-C**. For a given ROI, we calculated the average of the bone thickness measurements across the entire ROI (**Supplementary Fig. 2A**) and repeated this for every skull in the cohort (**Supplementary Fig. 2B**). The average thickness values were stored in vectors by cohort for analysis (**Supplementary Fig. 2C**).

### Validation of μCT scan measurements

Full thickness burr holes were drilled into the parietal bones of three half skulls during acute surgeries. The skulls were scanned using both the μCT scanner and a custom-built optical coherence tomography (OCT) scanner with a center wavelength of 1300 nm and a resolution of 7 μm. The thickness of the bone adjacent to each burr hole was measured with both scan modalities and the results were compared. We found that the thickness measurements agreed to within ± 40 μm. Given the μCT resolution of 21 μm, the disagreement is reasonably minor. This experiment confirmed that the μCT scans and pipeline accurately measured the skull bone thickness.

## SOFTWARE AVAILABILITY STATEMENT

The MATLAB software for μCT processing and analysis is available at our GitHub repository: www.github.com/bsbrl.

## ACKNOWLEDGEMENTS

SBK acknowledges funds from the National Institutes of Health (NIH) Brain Initiative Award 1RF1NS113287 and National Institute of Neurological Disorders and Stroke (NINDS) Award R01NS111028. μCT scans were performed at the Minnesota Dental Research Center for Biomaterials and Biomechanics with the assistance of Bonita VanHeel. OCT scans were performed in Dr. Taner Akkin’s lab with the assistance of Tianqi Li. We thank Skylar Fausner for assistance with animal experiments. We would also like to thank Kapil Saxena and Travis Beckerle for useful comments on the manuscript.

## AUTHOR CONTRIBUTIONS

B. R. G. and Z. S. N. prepared samples for scanning. B. R. G. performed the μCT scans and scan reconstruction. B. R. G. developed the MATLAB-based scan processing and analysis software. B. R. G. and Z. S. N. validated the skull thickness measurements. B. R. G. and S. B. K. wrote the manuscript.

## INSTITUTIONAL APPROVAL

All animal experiments described in this paper were approved by the University of Minnesota’s Institutional Animal Care and Use Committee (IACUC).

## DATA AVAILABILITY STATEMENT

All data included in this manuscript will be made available by the authors upon reasonable request.

## COMPETING INTERESTS

The authors declare no competing interests.

## LIST OF SUPPLEMENTARY MATERIAL

**Supplementary Figure 1**

Figure illustrating the registration of skulls to a common reference to create an average skull for each cohort.

**Supplementary Figure 2**

Figure showing quantification of average dorsal bone thickness differences between cohorts.

**Supplementary Figure 1.**
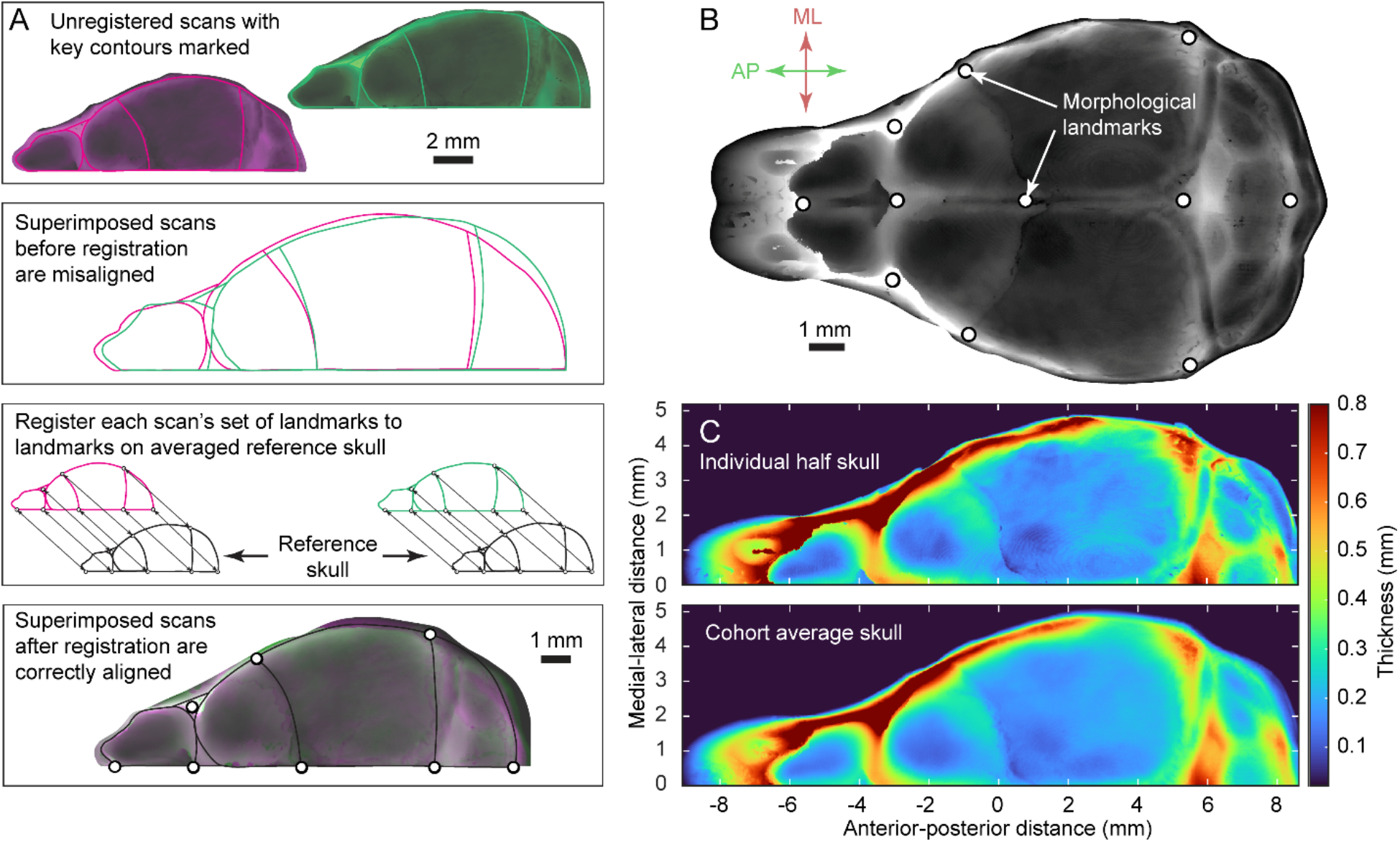
Registering skulls to common reference to create average skull for each cohort: **(A)** Method for registering skulls. Cranial form varied between skulls, so registration was necessary before averaging. Using a control point registration method from the MATLAB Image Processing Toolbox, we manually selected landmarks on each skull. Each measurement matrix was then warped to fit its landmarks to a set of averaged reference control points using a piecewise linear transformation method. **(B)** The control points used during registration, selected based on the morphological landmarks which were consistently identifiable in scans. **(C)** Comparing pseudo-color plots of dorsal skull bone thickness in (top) an individual skull and (bottom) the corresponding cohort average registered skull. Similarities in bone thickness structure between individual skulls and the cohort average skull indicate successful registration.

**Supplementary Figure 2.**
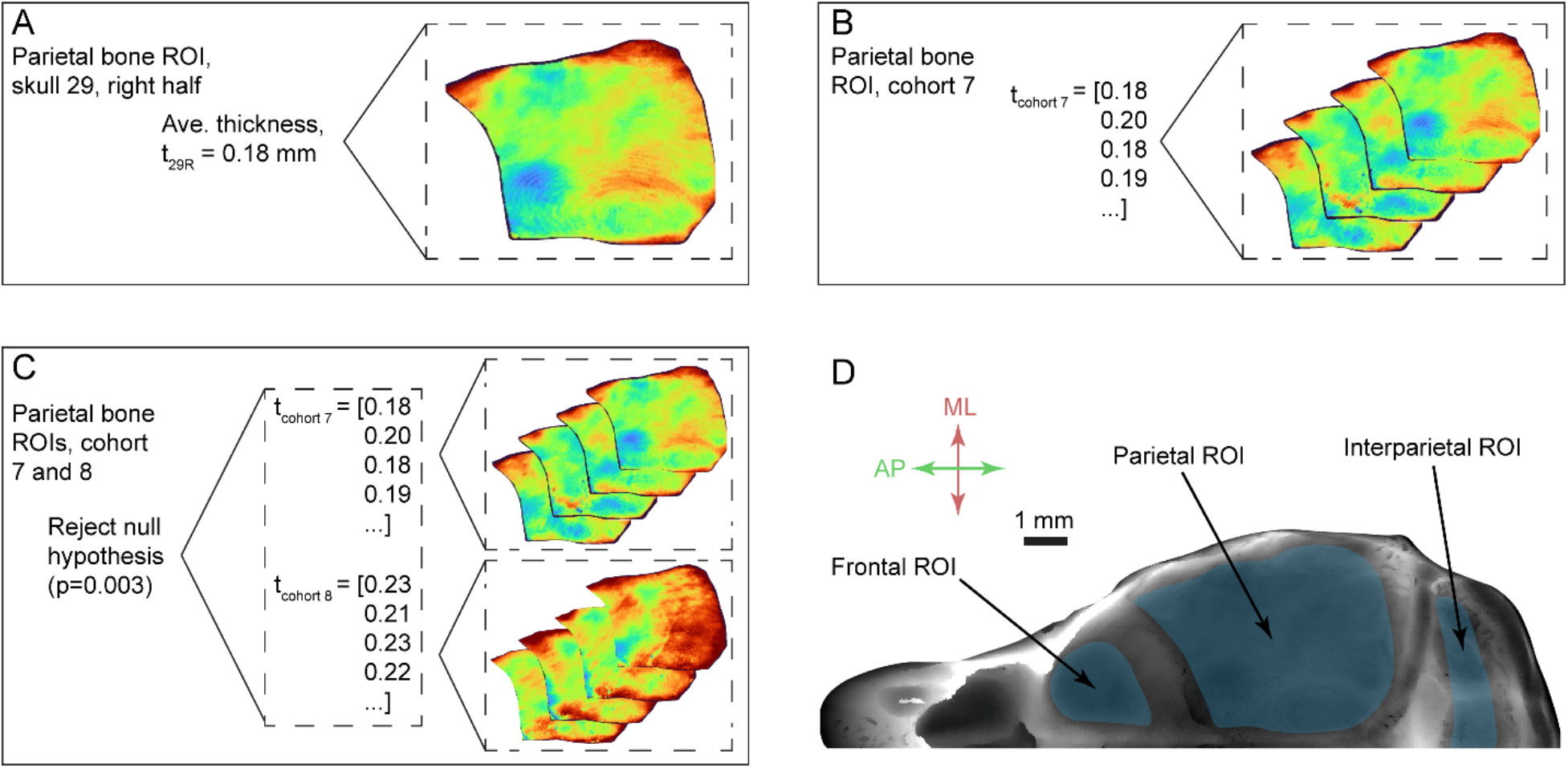
Method for quantifying average dorsal bone thickness differences between cohorts: **(A)** For a given ROI, the average of the bone thickness values was calculated for each half skull in the cohort. **(B)** Each average bone thickness metric from all the half skull ROIs in the cohort was stored in a vector. **(C)** A Students’ two-sample t-test was performed on the average thickness vectors from each set of two cohorts with only one variable differing between them. **(D)** Locations of ROIs: frontal, parietal, and interparietal bone ROIs, avoiding sutures.

## Notes

### Competing Interest Statement

The authors have declared no competing interest.

## REFERENCES

[1] Ellenbroek B. & Youn J. Rodent models in neuroscience research: is it a rat race? Dis. Model Mech. 9, 10 1079–1087 (2016). https://doi.org/10.1242/dmm.026120

[2] Madisen L. et al. Transgenic Mice for Intersectional Targeting of Neural Sensors and Effectors with High Specificity and Performance. Neuron 85, 5 942–958 (2015). https://doi.org/10.1016/j.neuron.2015.02.022

[3] Daigle T. L. et al. A Suite of Transgenic Driver and Reporter Mouse Lines with Enhanced Brain-Cell-Type Targeting and Functionality. Cell 174, 2 465–480 (2018). https://doi.org/10.1016/j.cell.2018.06.035

[4] Vora, S. R., Camci, E. D. & Cox, T. C. Postnatal Ontogeny of the Cranial Base and Craniofacial Skeleton in Male C57BL/6J Mice: A Reference Standard for Quantitative Analysis. Front. Physiol. 6, 417 (2016). https://doi.org/10.3389/fphys.2015.00417

[5] Kawakami, M. & Yamamura, K. I. Cranial bone morphometric study among mouse strains. BMC Evol. Biol. 8, 73 (2008). https://doi.org/10.1186/1471-2148-8-73

[6] Matthaei K. I. Genetically manipulated mice: a powerful tool with unsuspected caveats. J. Physiol. 582, 2 481–488 (2007). https://doi.org/10.1113/jphysiol.2007.134908

[7] Dana H. et al. Thy1-GCaMP6 Transgenic Mice for Neuronal Population Imaging In Vivo. PLoS One 9, 9 e108697 (2014). https://doi.org/10.1371/journal.pone.0108697

[8] Pak, N. et al. Closed-loop, ultraprecise, automated craniotomies. J. Neurophysiol. 113, 10 3943–3953 (2015). https://doi.org/10.1152/jn.01055.2014

[9] Rynes, M.L. et al. Assembly and operation of an open-source, computer numerical controlled (CNC) robot for performing cranial microsurgical procedures. Nat. Protoc. 15, 1992–2023 (2020). https://doi.org/10.1038/s41596-020-0318-4

[10] Ghanbari, L. et al. Craniobot: A computer numerical controlled robot for cranial microsurgeries. Sci. Rep. 9, 1023 (2019). https://doi.org/10.1038/s41598-018-37073-w

[11] Franklin, K. B. J. & Paxinos, G. Paxinos and Franklin’s the Mouse Brain in Stereotaxic Coordinates. Fourth ed. Amsterdam: Academic Press an imprint of Elsevier; 2013.

[12] Allen Institute for Brain Science (2011). Allen Reference Atlas - Mouse Brain [brain atlas]. https://atlas.brain-map.org

[13] Holtmaat, A. et al. Long-term, high-resolution imaging in the mouse neocortex through a chronic cranial window. Nat. Protoc. 4, 1128–1144 (2009). https://doi.org/10.1038/nprot.2009.89

[14] Ghanbari, L. et al. Cortex-wide neural interfacing via transparent polymer skulls. Nat. Commun. 10, 1500 (2019). https://doi.org/10.1038/s41467-019-09488-0

[15] Hong, G. et al. Through-skull fluorescence imaging of the brain in a new near-infrared window. Nature Photon 8, 723–730 (2014). https://doi.org/10.1038/nphoton.2014.166

[16] Jo Y. et al. Through-skull brain imaging in vivo at visible wavelengths via dimensionality reduction adaptive-optical microscopy. Sci. Adv. 8, 30 (2022). https://doi.org/10.1126/sciadv.abo4366.

[17] Szu, J. I. et al. Thinned-skull Cortical Window Technique for In Vivo Optical Coherence Tomography Imaging. J. Vis. Exp. 69, e50053 (2012). https://doi.org/10.3791/50053

[18] Wang, T. et al. Three-photon imaging of mouse brain structure and function through the intact skull. Nat. Methods 15, 789–792 (2018). https://doi.org/10.1038/s41592-018-0115-y

[19] Drew, P. et al. Chronic optical access through a polished and reinforced thinned skull. Nat. Methods 7, 981–984 (2010). https://doi.org/10.1038/nmeth.1530

[20] Nsiangani, A. et al. Optimizing intact skull intrinsic signal imaging for subsequent targeted electrophysiology across mouse visual cortex. Sci. Rep. 12, 2063 (2022). https://doi.org/10.1038/s41598-022-05932-2

[21] Silasi, G., Xiao, D., Vanni, M. P., Chen, A. C. N. & Murphy, T. H. Intact skull chronic windows for mesoscopic wide-field imaging in awake mice. J. Neurosci. Methods 267, 141–149 (2016). https://doi.org/10.1016/j.jneumeth.2016.04.012

[22] Rindone, A. N. et al. Quantitative 3D imaging of the cranial microvascular environment at single-cell resolution. Nat. Commun. 12, 6219 (2021). https://doi.org/10.1038/s41467-021-26455-w

[23] Otsu, N. A Threshold Selection Method from Gray-Level Histograms. IEEE Trans. Syst. Man Cybern. 9, 1 62–66 (1979). https://doi.org/10.1109/TSMC.1979.4310076

